# Ultrasound-Triggered Chemotherapy Extends Survival in a Genetically Engineered Glioblastoma Model

**DOI:** 10.64898/2026.06.29.735435

**Authors:** Joshua Antonio Whiting, Audri Yasmin Al Hasan Dara, James Francis Kwan, Anna Edmunds, Sheri Holmen, Jan Kubanek

## Abstract

Glioblastoma (GBM) remains one of the most lethal primary brain tumors, in part because the blood–brain barrier (BBB), restricts delivery of most systemically administered chemotherapeutics. Although focused ultrasound (fUS) can transiently increase BBB permeability, therapeutic efficacy remains limited by reliance on systemic drug exposure and heterogeneous intratumoral distribution. Here, we report a pressure-gated ultrasound-triggered drug delivery strategy that enables localized intravascular release of chemotherapy at the site of sonication. Freebase doxorubicin and afatinib were encapsulated within ultrasound-sensitive mPEG–PDLLA/PFOB microdroplets and administered systemically to N-TVA::Ink4a/Arf^lox/lox^;Pten^lox/lox^ mice bearing genetically engineered glioblastomas. Animals received repeated transcranial focused ultrasound over a 30-day treatment period. Ultrasound-triggered release of the dual-drug formulation significantly extended survival compared with untreated controls, with median survival increased by over two weeks – approximately a 30% improvement. Furthermore, this survival improvement was reflected in histological analysis, showing decreased tumor burden and severity. These improvements were not found in any control groups, demonstrating that spatially and temporally controlled intravascular drug release can substantially improve therapeutic efficacy in an aggressive immunocompetent glioblastoma model. These findings support pressure-gated ultrasound-triggered chemotherapy as a promising activation-based strategy for overcoming BBB-associated delivery limitations and improving outcomes in malignant brain tumors.

**Graphical Abstract:** 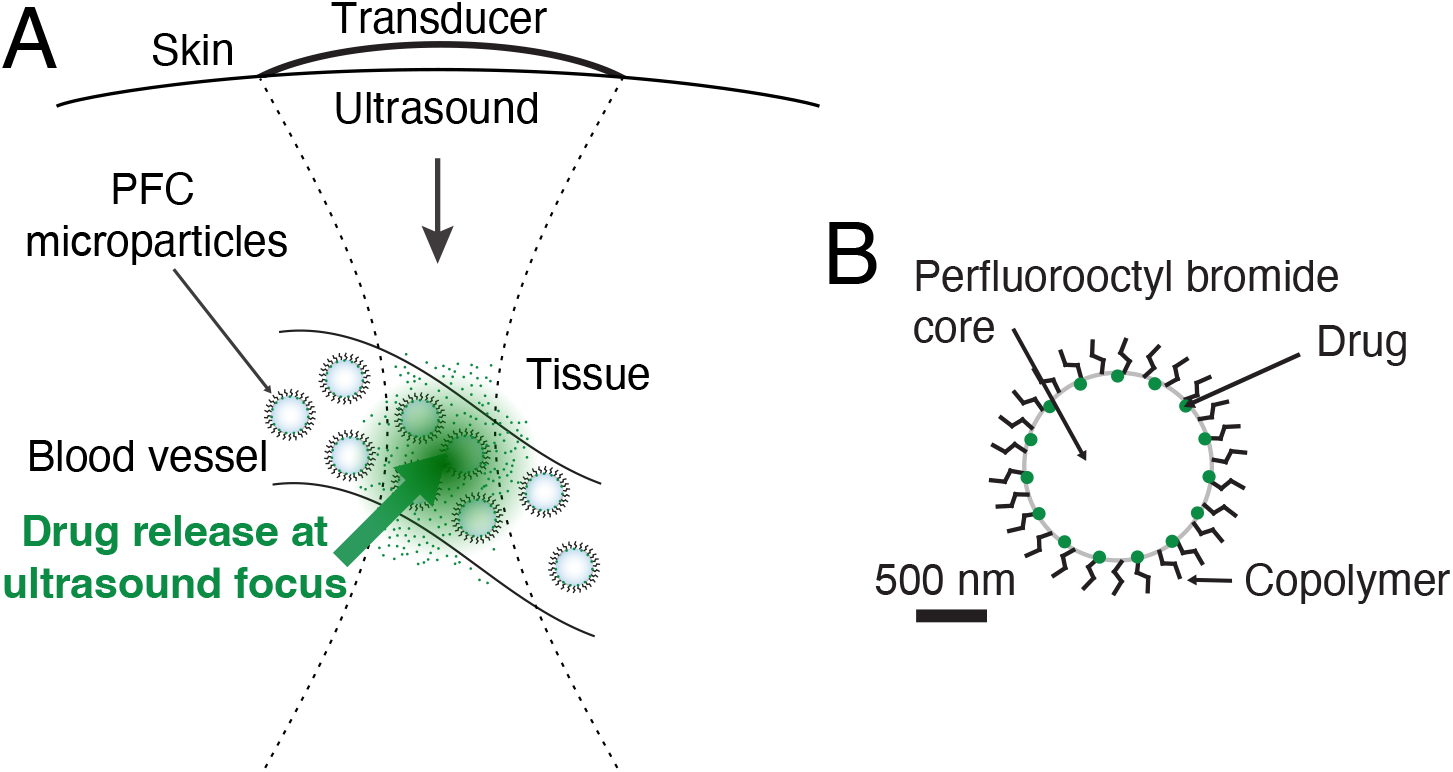

**Highlights:**

- Pressure-gated focused ultrasound enables localized release of doxorubicin and afatinib in glioblastoma.
- Ultrasound-triggered chemotherapy significantly extends survival in a genetically engineered immunocompetent GBM model.
- Local activation outperforms systemic administration of identical drug combinations.
- This strategy shifts focused ultrasound therapy from general BBB opening to spatially controlled drug activation.

## 1. Introduction

Glioblastoma multiforme (GBM) is the most aggressive and lethal primary brain tumor [1]. Current standard-of-care treatment, consisting of maximal surgical resection followed by radiotherapy and temozolomide chemotherapy, provides only a modest survival benefit, with median survival remaining approximately 15 months following diagnosis [1, 2]. This poor prognosis is driven by rapid tumor proliferation, diffuse infiltration into surrounding brain tissue, extensive molecular heterogeneity, and the development of therapeutic resistance.

A major obstacle to effective GBM treatment is the blood–brain barrier (BBB), a highly selective endothelial interface that restricts the passage of most systemically administered therapeutics into the brain parenchyma [3, 4]. Consequently, many potent antineoplastic agents fail to achieve therapeutic concentrations within brain tumors despite demonstrating substantial efficacy in vitro. Increasing systemic dosing to compensate for poor brain penetration often results in dose-limiting toxicities that prevent clinical implementation.

Among the agents limited by BBB penetration are doxorubicin and afatinib. Doxorubicin is a broad-spectrum anthracycline chemotherapeutic that induces DNA damage and apoptosis through inhibition of topoisomerase II, while afa-tinib is an irreversible epidermal growth factor receptor (EGFR) inhibitor that suppresses signaling pathways frequently dysregulated in GBM [2, 5, 6, 7, 8]. Although both agents demonstrate antitumor activity, their clinical utility in brain tumors is limited by poor BBB permeability, rapid systemic clearance, and significant off-target toxicity.

Focused ultrasound (fUS) has emerged as a promising technology for targeted drug delivery to the brain. Most current approaches employ ultrasound-mediated BBB disruption, often in conjunction with circulating microbubbles, to transiently increase vascular permeability and facilitate drug entry into brain tissue [3, 4, 9, 10]. While this strategy can enhance drug penetration, therapeutic efficacy remains dependent upon systemic drug circulation, diffusion through heterogeneous tumor tissue, and maintenance of sufficient drug concentrations after BBB opening.

An alternative strategy is ultrasound-triggered intravascular drug release. Rather than relying on passive diffusion after BBB disruption, therapeutic agents are encapsulated within ultrasound-sensitive carriers and released directly within the tumor vasculature at the site and time of ultrasound exposure. This approach enables spatially and temporally controlled drug delivery, generating high local drug concentrations while limiting systemic toxicity [11, 12, 13, 14, 15, 16].

Polymeric and lipid-based ultrasound-responsive carriers have demonstrated efficient encapsulation and controlled release of chemotherapeutics including doxorubicin, paclitaxel, and docetaxel [13, 14, 15, 16]. In particular, fluorocarbon-containing systems enable pressure-gated release under clinically relevant ultrasound parameters [17, 18].

Our group previously developed an ultrasound-sensitive drug delivery platform based on methoxy poly(ethylene glycol)-poly(D,L-lactide) (mPEG-PDLLA) copolymer microdroplets containing perfluorooctyl bromide (PFOB), demonstrating remotely controlled release in deep brain regions of non-human primates with excellent safety and repeatability [19, 20, 21, 17]. More recently, we adapted this platform to encapsulate hydrophobic chemotherapeutic agents, including doxorubicin and afatinib, while preserving pressure-dependent ultrasound-triggered release characteristics [22, 23, 24].

In the present study, we evaluated whether pressure-gated ultrasound-triggered release of doxorubicin and afatinib could improve therapeutic outcomes in a genetically engineered (N-TVA::Ink4a/Arf^lox/lox^;Pten^lox/lox^) glioblastoma mouse model, as previously characterized by Shin et al. [25]. We hypothesized that localized intravascular release would increase tumor drug exposure while reducing systemic toxicity, resulting in prolonged survival and favorable tumor biological changes.

## 2. Materials and Methods

### 2.1. Animal Model and Treatment Groups

All animal procedures were approved by the University of Utah Institutional Animal Care and Use Committee (IACUC) and conducted in accordance with institutional guidelines.

Glioblastoma tumors were generated in N-TVA;Ink4a/Arf^lox/lox^;Pten^lox/lox^ mice using the RCAS/tv-a system as previously described by Shin et al. [25]. Briefly, immunocompetent newborn mice of both sexes received intracranial injections of DF-1 producer cells carrying RCAS-HB-EGF and RCAS-Cre constructs, resulting in somatic deletion of *Ink4a/Arf* and *Pten* and induction of high-grade gliomas. This genetically engineered model recapitulates key histopathological and molecular features of human glioblastoma and has been shown to exhibit elevated EGFR signaling and gene expression consistent with the classical GBM subtype [25].

Treatment was initiated at weaning (approximately postnatal day 21), a time point at which tumors are reliably established in this model [25], and littermates were distributed across treatment groups when possible. Treatment groups consisted of: (1) focused ultrasound (fUS) with dual-drug microdroplets containing doxorubicin and afatinib (*n* = 12); (2) fUS with doxorubicin-loaded microdroplets (*n* = 13); (3) fUS with afatinib-loaded microdroplets (*n* = 13); (4) fUS with empty microdroplets (*n* = 12); (5) systemic doxorubicin (*n* = 13); (6) systemic afatinib (*n* = 13); (7) systemic doxorubicin and afatinib combination therapy (*n* = 13); and (8) untreated controls (*n* = 16).

Animals were monitored daily for neurological deficits, body weight changes, and overall health status. Mice reaching predefined humane endpoints, including severe neurological impairment, inability to access food or water, or greater than 20% body weight loss, were euthanized. Survival was defined as the time from birth to euthanasia due to humane endpoint criteria.

### 2.2. Microdroplet Synthesis

Ultrasound-sensitive microdroplets were synthesized using a mPEG-PDLLA (2,000:2,200 g/mol) diblock copolymer shell and a perfluorooctyl bromide (PFOB) core [22]. Drug encapsulation was achieved using a solvent evaporation and emulsification process optimized for hydrophobic chemotherapeutics.

Briefly, doxorubicin free base or afatinib dissolved in DMSO was combined with polymer dissolved in tetrahydrofuran (THF), followed by solvent evaporation to form a thin film. The film was rehydrated in phosphate-buffered saline and emulsified with PFOB using probe sonication to form stable microdroplets. Unencapsulated drug and excess polymer were removed via repeated centrifugation and washing.

Previously characterized formulations exhibited mean diameters of 1-2 µm, encapsulation efficiencies of 40–47%, and pressure-dependent release with maximal activation at 1.3 MPa and above[22]. Batches were prepared the morning of treatments, stored at 4^?^C until use, and remained on ice throughout use.

### 2.3. Focused Ultrasound Treatment

Animals received retro-orbital injections of either microdroplet formulations or systemic drug controls at equivalent drug dosages. Focused ultrasound was applied transcranially using a 300 kHz H-115 transducer (Soniq Concepts) coupled with a gel-filled focusing cone and ultrasound gel, with the focal area enveloping the brain region as characterized in water-bath experiments.

Sonication consisted of 10-ms pulses at 10 Hz for 60 seconds with a duty cycle of 10%, repeated three times per week over a 30-day treatment period. This regimen was selected based on prior optimization studies demonstrating pressure-dependent release without tissue damage [18, 17].

### 2.4. Histology and Immunohistochemistry

Ex vivo Brains were collected in formalin and fixed for 24–36 hours, then stored in 70% isopropyl alcohol. At study completion, all brains were batch processed, paraffin-embedded, and sectioned at 4 µm for histological and immunohistochemical analysis. Sections were stained for hematoxylin and eosin (H&E), Ki-67, GFAP, p-ERK, and cleaved caspase-3 (CC3). H&E slides were used to score tumors on severity based on neuropathological indicators such as pseudopalisading necrosis, aberrant vasculature, and additional histopathological features relevant to glioblastoma grade. Ki-67 was selected as a marker of tumor proliferation. GFAP was selected to confirm glial lineage of identified tumors. p-ERK was selected as a marker of afatinib action because it is downstream of the EGFR pathway inhibited by afatinib. Finally, CC3 was selected as a marker of apoptosis, thereby serving as a marker of doxorubicin effect, which induces double-stranded DNA breaks and apoptosis.

### 2.5. Statistical Analysis

Kaplan–Meier survival curves were compared using the Mantel–Cox log-rank test. Histological tumor grading was performed blinded and each slide was assigned a tumor score of 0-4, with 0 indicating no signs of tumor burden and 4 indicating severe disease. Quantitative IHC data will be analyzed using Student’s t-test or Mann–Whitney U test, dependent upon data normality. For all statistical tests, a p-value <0.05 will be considered statistically significant.

## 3. Results

### 3.1. Survival Benefit of Ultrasound-Triggered Chemotherapy

Mice receiving fUS-triggered release of dual-drug microdroplets exhibited a median survival of 63.5 days compared with 49 days in untreated controls, representing an approximately 30% increase in survival (p= 0.02). This improvement was not observed with systemic administration of the same agents, single-agent with fUS administration, or unloaded microdroplets with fUS.

**Figure 1:**
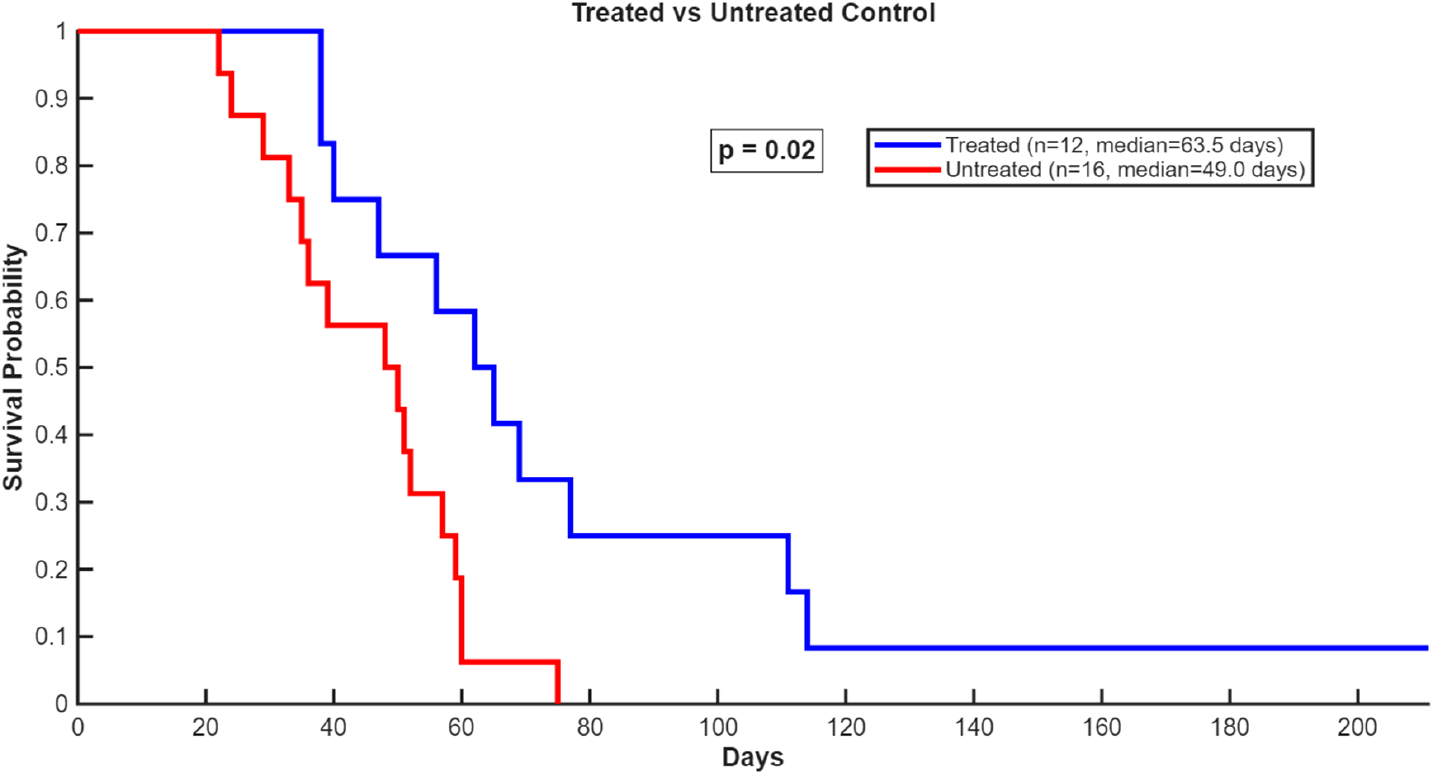
Kaplan–Meier survival analysis.

### 3.2. Histological And Immunohistochemical Analysis

Histological analysis demonstrated reduced tumor burden and lower pathological grade in animals receiving ultrasound-triggered combination therapy, as shown in Figure 2. Additionally, the only animal showing no signs of GBM at study conclusion was in the dual-drug microdroplet + fUS treatment group. Quantitative image analysis of immunohistochemical markers is ongoing and will be reported in a future version of the manuscript.

**Figure 2:**
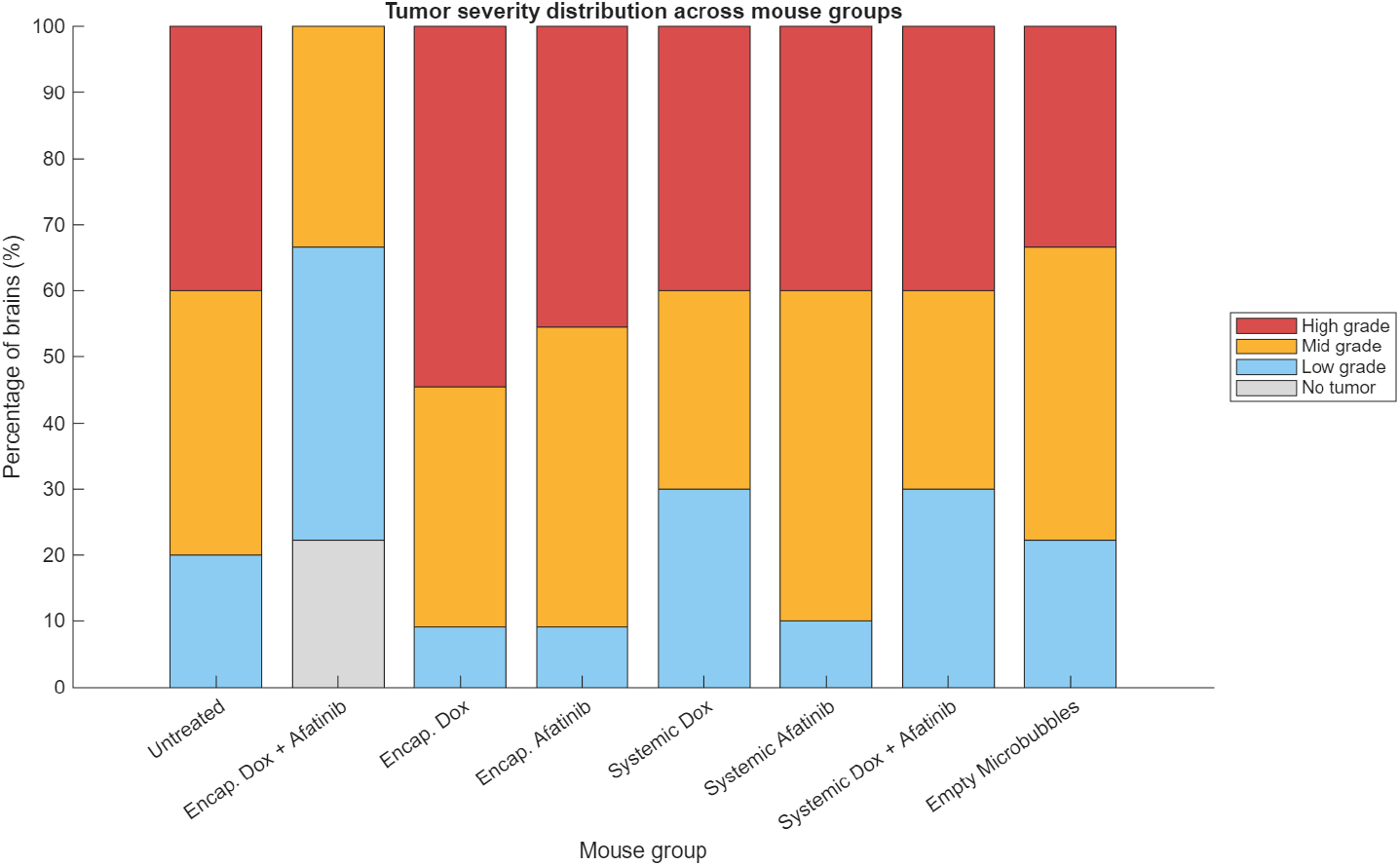
Representative tumor grading results.

## 4. Discussion

Focused ultrasound-mediated drug delivery has been widely investigated as a strategy to improve central nervous system drug exposure, most commonly through transient blood–brain barrier (BBB) disruption [3, 4, 9]. While this approach increases permeability to circulating therapeutics, its efficacy remains constrained by systemic pharmacokinetics, heterogeneous tumor perfusion, and limited spatial control over drug deposition.

In contrast, the present study demonstrates a fundamentally different delivery paradigm: pressure-gated, ultrasound-triggered intravascular drug release. Rather than enhancing passive transport across the BBB, this strategy enables direct acoustic activation of drug release within tumor-associated vasculature, producing a localized, high-concentration pharmacologic bolus precisely at the site of ultrasound exposure. This mechanism decouples therapeutic efficacy from systemic circulation time and BBB permeability.

Our findings demonstrate that ultrasound-triggered release of doxorubicin and afatinib significantly prolongs survival in an aggressive, immunocompetent genetically engineered glioblastoma model. Importantly, equivalent systemic administration of the same drug combinations did not improve survival, indicating that therapeutic benefit arises from spatially confined activation rather than increased systemic exposure alone.

These results are consistent with prior studies demonstrating ultrasound-enhanced drug delivery and acoustic activation of nanocarriers for local chemotherapy [13, 11, 15, 14, 26]. However, most prior work has not demonstrated survival benefit in genetically engineered glioblastoma models, highlighting the translational relevance of the present findings.

This platform builds upon our previous demonstrations of safe and repeatable ultrasound-triggered release in deep brain structures using mPEG–PDLLA/ PFOB microdroplets in non-human primates [19, 20], as well as recent adaptations enabling efficient encapsulation of hydrophobic chemotherapeutics while preserving pressure sensitivity [22]. Together, these results support the broader concept of activation-based chemotherapy, in which therapeutic effect is governed by an external physical trigger rather than systemic pharmacokinetics.

While this study focused on glioblastoma and two chemotherapeutic agents, the underlying mechanism may be broadly applicable to other hydrophobic drugs and disease contexts where spatial control of drug activity is critical.

In conclusion, pressure-gated focused ultrasound enables activation-based intravascular chemotherapy that significantly improves survival in a genetically engineered glioblastoma model. This approach represents a shift from permeability-based delivery toward spatially controlled therapeutic activation and provides a promising platform for the treatment of malignant brain tumors.

## 5. Acknowledgements

The authors thank the members of the University of Utah’s Neuropathology department for assistance with IHC analysis and neuropathology, and the members of the OneTarget Lab for helpful discussions and technical assistance. This work was supported by National Science Foundation (Grant No. CBET 2325125), the National Institutes of Health (Grant No. R61/R33), and the Huntsman Mental Health Institute and College of Engineering at the University of Utah. The content is solely the responsibility of the authors and does not necessarily represent the official views of the funding agencies.

## 6. Author contributions

Joshua Antonio Whiting: Conceptualization, investigation, methodology, formal analysis, visualization, writing – original draft.

Audri Yasmin Al Hasan Dara: Investigation, methodology, writing – review and editing.

James Kwan: Investigation, writing – review and editing.

Anna Edmunds: Investigation and methodology.

Sheri Holmen: Conceptualization, supervision, resources, writing – review and editing.

Jan Kubanek: Conceptualization, supervision, resources, funding acquisition, writing – review and editing.

